# Characterization of a membrane enzymatic complex for heterologous production of poly-γ-glutamate in *E. coli*

**DOI:** 10.1101/2020.07.17.209205

**Authors:** Bruno Motta Nascimento, Nikhil U. Nair

**Affiliations:** Department of Chemical and Biological Engineering, Tufts University, Medford, MA, 02155, USA

**Author notes:** Present/permanent address: Science and Technology Center #276, 4 Colby St., Medford, MA 02155.

**Keywords:** heterologous, poly-gamma-glutamate, synthetase, biopolymer, localization, membrane

## Abstract

Poly-γ-glutamic acid (PGA) produced by many *Bacillus* species is a polymer with many distinct and desirable characteristics. However, the multi-subunit enzymatic complex responsible for its synthesis, PGA Synthetase (PGS), has not been well characterized yet, in native nor in recombinant contexts. Elucidating structural and functional properties are crucial for future engineering efforts aimed at altering the catalytic properties of this enzyme. This study focuses on expressing the enzyme heterologously in the *Escherichia coli* membrane and characterizing localization, orientation, and activity of this heterooligomeric enzyme complex. In *E. coli*, we were able to produce high molecular weight PGA polymers with minimal degradation at titers of approximately 13 mg/L in deep-well microtiter batch cultures. Using fusion proteins, we observed, for the first time, the association and orientation of the different subunits with the inner cell membrane. These results elucidate provide fundamental structural information on this poorly studied enzyme complex and will aid future fundamental studies and engineering efforts.

**HIGHLIGHTS:**

- Successfully expressed active poly-γ-glutamate synthetase (PGS) in *E. coli*.
- Confirmed PGS localization at inner membrane of *E. coli*.
- Elucidated topology of PGS components in *E. coli* membrane.
- Culture and expression in microplates might allow future screening of a high number of samples.
- Faster production of poly-γ-glutamate in *E. coli* supernatant compared to *B. subtilis*.

## 1. INTRODUCTION

Poly-γ-glutamate synthetase (PGS) is a multimeric enzyme present in some *Bacillus* species and a few other organisms. While some other amide ligases are able to synthesize poly-glutamate with α-amide linkages between the amino and α-carboxyl groups of L-glutamate (Hamano et al., 2013; Kino et al., 2011), PGS produces an unusual anionic polymer with γ-amide linkages, where the amino group reacts with the side chain carboxyl group of glutamate. In its natural cell environment, this poly-γ-glutamate (PGA) polymer functions as a glutamate storage, a cryoprotective material, and a protection against protease attacks and pH changes near the cell surface (Ashiuchi and Misono, 2003, 2002). *Bacillus* cultures that produce PGA usually have a highly viscous appearance and they have been used in Japan for a long time to produce fermented food products such as natto (Ashiuchi and Misono, 2002). Each species/strain presents a different preference for glutamate enantiomer utilization and molecular size distribution of the final polymer. *Bacillus anthracis* has a PGA composed of only D-glutamate, while *Bacillus halodurans* uses only L-glutamate for polymer production.

For industrial applications, most studies have relied on biosynthesizing PGA in *B. subtilis* (Ashiuchi, 2013; Halmschlag et al., 2019; Park et al., 2005; Wang et al., 2017) and other *Bacillus* species (Feng et al., 2017, 2015; Ogunleye et al., 2015; Tian et al., 2014; Xavier et al., 2019; Yoon et al., 2000), although there have been significant efforts in moving the complex to recombinant hosts such as *E. coli* (Ashiuchi et al., 1999; Cao et al., 2013, 2011; Jiang et al., 2006; Liu et al., 2019; Wang et al., 2011), *Corynebacterium glutamicum* (Xu et al., 2019), and even tobacco plants (Tarui et al., 2005). A major advantage of recombinant hosts is that PGA synthesis can be decoupled from native cellular regulatory processes and engineered for higher productivity, yield, stereochemical composition, and molecular weight. Further, the absence of any endogenous PGA hydrolytic enzyme (e.g. PgsD and GGT (Ojima et al., 2019; Scoffone et al., 2013)) ensures higher product stability during culture. Despite significant molecular and bioprocess engineering efforts, there have been few efforts focused on characterizing the structure, assembly, and function of this enzyme complex. Current research findings support the hypothesis that PGS is a membrane-associated protein comprised of four subunits – PgsB, PgsC, PgsA, and PgsE, as proposed previously (Ashiuchi, 2013) – and that polymerization of glutamate occurs concurrently with secretion of the growing chain. However, the role and membrane localization of the different subunits have not been well-characterized. Based on all these previous insights, our study is focused on producing the PgsBCAE enzymatic complex from *B. subtilis* in *E. coli* and analyzing if it was enzymatically active and localized correctly to the membrane in this non-native host. We characterize PGS by co-expressing different combinations of its four subunits and fusing different reporters to assess membrane (co-)localization and orientation. We determine the orientation of the different subunits within the cytoplasmic and periplasmic space. Results from this work shed light into the assembly of PGS and will aid in future engineering efforts.

## 2. MATERIALS AND METHODS

### 2.1. Strains and culture media

Bacterial strains used for this project are listed in Table S1. *Escherichia coli* NEB5α competent strain was used during the cloning steps. *E. coli* MG1655(DE3)^ΔrecA,ΔendA^ was used for recombinant gene expression. Chemically competent cells were prepared by the calcium chloride/MES method. Luria-Bertani (LB) media (VWR Life Science, #97064-110) was used for propagation, preservation of bacterial cells and expression experiments. In the case of solid medium, 18 g/L of bacteriological agar was added, and whenever the cells where transformed with plasmids, the appropriate antibiotic was added to the cooled medium at the recommended final concentration (25 μg/mL chloramphenicol or 100 μg/mL ampicillin). Isopropyl β-D-1-thiogalactopyranoside (IPTG) was added for induction, and its concentration varied among experiments. Growth in liquid media was done in orbital shaker at 250 rpm and 37 °C, unless noted otherwise. “Magic” medium was used in initial expression trials, and constituted of 15 g/L LB broth (VWR Life Science, #97064-110), 5 g/L glucose, 10 g/L yeast extract, 2 g/L Tris, 4 mL/L glycerol, 55 mM K_2_HPO_4_, 15 mM KH_2_PO_4_, 10 mM MgSO_4_, and 10 mM (NH_4_)_2_SO_4_ prepared with tap water.

### 2.2. Cloning and expression

Plasmid extraction was performed using the E.Z.N.A^®^ Plasmid DNA Mini Kit I (OMEGA Bio-tek, #D6943-02). Horizontal DNA electrophoresis in agarose gel was performed in 1× TAE buffer according to (Sambrook and Russell, 2001). Purification of DNA fragments from agarose gels used the MicroElute^®^ Gel Extraction Kit (OMEGA Bio-tek, #D6294-02). DNA concentration was quantified by UV absorbance with the SpectraMax M3 microplate reader using a SpectraDrop Micro-Volume Microplate (Molecular Devices). DNA amplification with Taq DNA polymerase or Phusion^®^ High-Fidelity DNA polymerase was performed according to manufacturer recommendations (New England Biolabs, Thermo Fisher Scientific). DNA digestion with restriction enzymes, DNA ligation with T4 DNA ligase, and DNA assembly with NEBuilder kit were done according to the manufacturer recommendations (New England Biolabs). DNA sequences of constructs were confirmed by sequencing with the appropriate primers, which was performed by GENEWIZ (Boston Lab, Cambridge, MA). The *pgsBCAE* operon was amplified from *B. subtilis* 168 genome and used as template for all the constructs. In general, the PGS subunits were inserted in a pACYC-Duet1 plasmid under control of a T7 promoter. Different combinations of the subunits in a operon inducible by IPTG, generating the plasmids described on Table S2. Later, tagged versions of each subunit were created at the C terminus of each protein a 12 amino acid linker and different reporters (6xHis tag, sfGFP, mCherry, GFPuv, PhoA).

For construction of *E. coli* strain MGΔΔ-ΔphoA, the lambda red recombineering method was used (Datsenko and Wanner, 2000). In brief, MGΔΔ cells were first transformed with pKD46 for lambda red recombinase expression under an arabinose-induced promoter. A deletion cassette was constructed by amplifying part of pKD3 containing a *cat* gene flanked by FRT sites. Flanking both sides of the cassette it was added a 40 bp sequence homologous to the *phoA* gene. This cassette was electroporated in the MGΔΔ pKD46 cells for recombination at the locus site to occur. Selected colonies were finally transformed with pCP20 for removal of the chloramphenicol resistance by action of the FLP recombinase at the FRT sites. Deletion of *phoA* was confirmed by both sequencing of this genomic region and the absence of PhoA activity in the resulting cells.

### 2.3. PGA production

*E. coli* cells were grown in 10 mL culture tubes containing 5 mL of LB or rich media broth supplemented with 30 g/L L-glutamate, 10 μM IPTG, and 25 μg/mL chloramphenicol. Cultures were incubated at 30 °C, 250 rpm, for 24 h. *B. subtilis* strains were also grown in similar conditions, except without the addition of IPTG and antibiotic. The PGA produced in the supernatant was either measured directly or purified by a modified version of the copper precipitation method described by (Manocha and Margaritis, 2010; Yuan et al., 2019). In brief, 0.5 M CuSO_4_ was added to the supernatant and the solution was mixed by inversion and let stand for 1 h at room temperature. Tubes were centrifuged at 5,000 × g for 30 min and the pellets were later resuspended in phosphate buffered saline (PBS) pH 7.4 with 50 mM EDTA. The solution was dialyzed with SnakeSkin® Dialysis Tubing 3.5 kDa molecular weight cut-off (MWCO) (Thermo Fisher Scientific, #68035) to remove salts, followed by drying in a vacuum centrifuge (Eppendorf Vacufuge Concentrator 5301).

### 2.4. Analytical methods

#### 2.4.1. Polymer and amino acids detection

The concentration of dialyzed samples of PGA was performed by adapting the cetrimonium bromide (CTAB) turbidity method described by (Halmschlag et al., 2019). To every 100 μL of sample it was added 50 μL of CTAB solution (CTAB 0.1 M, NaCl 1 M) and incubated for 5 min without agitation. Turbidity of the resulting solution was measured at 400 nm. Based on a calibration curve using standards of pure PGA (Sigma-Aldrich, #G1049-100MG) in the concentration range of 0.25 – 0.025 g/mL, the concentration of γ-PGA from the culture supernatant was determined. PGA detection by dot blot with nylon membrane also uses a similar staining procedure. The membrane was briefly dried at 37 °C and 2.5 μL each sample was applied to the membrane. The loaded membrane was dried again at 37 °C for 30 min and fixed with 60 % ethanol for 20 min. Ethanol was evaporated at 37 °C and the dry membrane was stained with 0.04% methylene blue in methanol for 5 – 10 min. Final de-staining was done with 25 % ethanol for 30 min. PAGE separation of PGA was done in a NuPAGE Bis-Tris 4 – 12 % gel (Thermo Fisher Scientific, # NP0322BOX) and ran in 1× MOPS buffer (50 mM MOPS, 50 mM Tris, 0.1 % SDS, 1 mM EDTA, pH 7.7) at 120 V for 2 h. The gel was washed twice with distilled wash for 5 min and stained with 0.03 % methylene blue in 300 mM sodium acetate pH 5.2 for 15 min with a gentle rocker agitation. The gel was de-stained by multiple washes with deionized water until the background and clear against the PGA bands (Soto and Draper, 2012).

#### 2.4.2. PGA hydrolysis and HPLC

Purified PGA samples from *E. coli* were hydrolyzed 6 M HCl at 100 °C for 4 h in a vacuum sealed tube (Chemglass, #CG-4025-01). Acid was immediately removed with a vacuum centrifuge and pellet was resuspended in deionized water. The presence of glutamate in the hydrolyzed samples was analyzed on an Agilent 1100 Series HPLC System, equipped with a Poroshell 120 HILIC-Z column (guard: 2.5 × 5 mm, 2.7 μm; main: 2.1 × 150 mm, 2.7 μm) and a diode array detector (Agilent, G1315B) measuring a signal at 338 nm wavelength (bandwidth = 10 nm) using reference wavelength 390 nm (bandwidth = 20 nm). Amino acid samples were derivatized with Fluoraldehyde *o*-phthaldialdehyde reagent (Thermo Fisher Scientific, #26025) prior injection in the system. The flow rate was constant and set at 0.3 mL/min. Solvents were run in a gradient condition with mobile phase A consisting of 10 mM ammonium acetate at pH 9.0 in water and mobile phase B consisting of 10 mM ammonium acetate at pH 9.0 in water:acetonitrile 1:9. After sample injection, mobile phases were run according to the following conditions: 0 – 2 min: 0 % A – 100 % B; 2 – 15 min: linear gradient to 30 % A – 70 % B; 15 – 16 min: linear gradient to 45 % A – 55 % B; 16 – 20 min: linear gradient to 0 % A – 100 % B; 20 – 25 min: 0 % A – 100 % B.

#### 2.4.3. Immunodetection of subunits

Subunits of PGS with 6xHis tags were detected by western blot. Expressing cells were recovered from culture media and resuspended in 2500 μL phosphate buffered saline (PBS) with 50 μL lysozyme 5 mg/mL, 5 μL DNaseI 5 mg/mL, and 2 μL 1M phenylmethylsulfonyl fluoride (PMSF). The cell suspension was lysed by sonication in a BRANSON Sonifier 150 (10 s ON, 1 min OFF, 5 min total ON, Amplitude 40 % with microtip). Following sonication, the tubes were centrifuged 3,000 ×g for 15 min at 4 °C to remove debris and un-lysed cells. Proteins from this lysate was quantified by BCA method (Abelson and Simon, 2009) and diluted to same concentration to load similar amounts of protein in each gel lane. These samples were separated in a 4 – 12 % NuPAGE Bis-Tris gel and ran in 1× MES buffer at 120 V until loading dye reached the bottom of the gel. Protein staining was done with SimplyBlue SafeStain (Thermo Fisher Scientific, #LC6060) according to manufacturer instructions. Transference to a PVDF membrane for western blot was done with the Invitrogen Xcell II blot module. Membrane was blocked with 5 % skim milk and antibody labeling used a primary mouse monoclonal anti-6xHis (Thermo Fisher, #MA1-21315) and a secondary rabbit anti-mouse IgG with HRP (Abcam, #ab6728). Chemiluminescence was detected with SuperSignal West Dura Exteded Duration Substrate (Thermo Fisher Scientific, #34075).

#### 2.4.4. Microscopy of labeled subunits

Cells were pelleted and washed with 1× PBS solution for preparation of microscope slides. A thin 2 % (w/v) agarose pad was prepared on a glass slide and 1 μL of cell suspension was added on top and air dried for about 5 min before placing a glass cover slip (Ke et al., 2016). Imaging was performed with a DMi8 automated inverted microscope (Leica Microsystems, #11889113) equipped with a CCD camera (Leica Microsystems, #DFC300 G), and a TXR (Leica Microsystems, #11525310) and YFP (Leica Microsystems, #11525306) filter cube. The Z-stack of captured images was processed in a blind deconvolution to remove out of focus fluorescence with the LAS X software version 3.3.3.16958 (Leica Microsystems, #11640612) with 3D deconvolution package (Leica Microsystems, #11640865).

#### 2.4.5. Detection of membrane orientation

Fusions to PhoA (alkaline phosphatase) are the most widespread periplasmic reporter fusions. Colonies expressing periplasmic PhoA fusions can be visually screened by supplementation of the agar medium with a PhoA-specific substrate 5-bromo-4-chloro-3-indolyl phosphate (X-Pho), yielding blue colonies when enzyme is active in the periplasm (Karimova and Ladant, 2017). Cells were streaked in a LB plate supplemented with 100 μg/mL X-Pho and grown at 30 °C for 4 h. After this period, a filter disc containing 5 nmol IPTG was plated at the center and the cells continued to grow at 30 °C for 24 h. Plates were then photographed to check development of a blue color due to PhoA activity. Conversely, GFPuv folds efficiently in the cytoplasm but does not form stable structure when targeted to the periplasm by a Sec-type signal peptide. GFPuv does fold properly, however, when attached to cytoplasmic domains of inner-membrane proteins (Drew et al., 2002). Cells were inoculated in 5mL LB-IPTG 10 μM media and induced at 30 °C for 4 h with shaking at 250 rpm. After this period, OD_600_ and fluorescence (excitation: 395 nm, emission: 509 nm) were measured with black microplates plates with clear bottom in a SpectraMax M3 microplate reader.

### 2.5. Bioinformatic analysis

An initial prediction of each protein interaction with the cell membrane was performed with the Constrained Consensus TOPology prediction server (CCTOP) (Dobson et al., 2015). It uses a combination of prediction methods (HMMTOP, Memsat, Octopus, Philius, Phobius, Pro, Prodiv, Scampi-single, Scampi-msa, TMHMM, SignalP) and generates a consensus topology with increased prediction accuracy. Homology model were also constructed with the SWISS-MODEL server (Schwede et al., 2003).

## 3. RESULTS AND DISCUSSION

### 3.1. Functional expression of recombinant PGA synthetase in E. coli

Initially, *E. coli* MGΔΔ cells expressing all PGS subunits (PgsBCAE) and wild type *B. subtilis* strains (168 and 6633) were cultured in “Magic” medium supplemented with 30 g/L L-glutamate. Cultures were grown at 30 °C (*E. coli*) and 37 °C (*B. subtilis*) – a lower incubation temperature was chosen for the former because burden of expression resulted in poor growth, probably due to membrane destabilization by the enzyme complex and the polymer produced. The crude fermented broth was recovered and run on a PAGE and the PGA was detected using methylene blue staining (Fig 1A). We found that the polymer produced by these bacteria have a high molecular weight (HMW), comparable to the pure commercial HMW PGA (>750 kDa). To reduce burden of expression, we moved the operon to a low-copy vector, which significantly improved growth of the *E. coli* transformants, allowing us to switch from “Magic” medium to glutamate-supplemented LB. We then tested which of the components are essential for PGA synthesis using strains expressing PgsBCAE, PgsBCA, PgsBCE, PgsBC, PgsBA, PgsB, or PgsA. Cultivation conditions were scaled down to a 24-deepwell plate containing 5 mL of media, sealed with aluminum film, and agitated at 700 rpm in a microplate shaker. After 24 h, a fraction of the supernatant was collected and precipitated with a copper solution as described in materials and methods to obtain a clearer PAGE staining result (Figure 1B). With purified PGA, we could also quantify the polymer by the CTAB method and estimate a concentration of 13 mg/L and < 1.6 mg/L by PgsBCAE and PgsBCA expressing *E. coli*, respectively (Figure S1). None of the other constructs tested in *E. coli*, nor wildtype *B. subtilis* 168, produced any detectable amounts of PGA under these conditions. These results also indicate that while PgsE is not essential for activity or processivity (Ashiuchi et al., 2013; Yamashiro et al., 2011a, 2011b), its presence enhances PGS activity. The non-essentiality of PgsE corroborates with other published studies where only the PgsBCA subunits were used for expression (Ashiuchi and Misono, 2002; Cao et al., 2011, 2010; Jiang et al., 2006).

**Figure 1.**
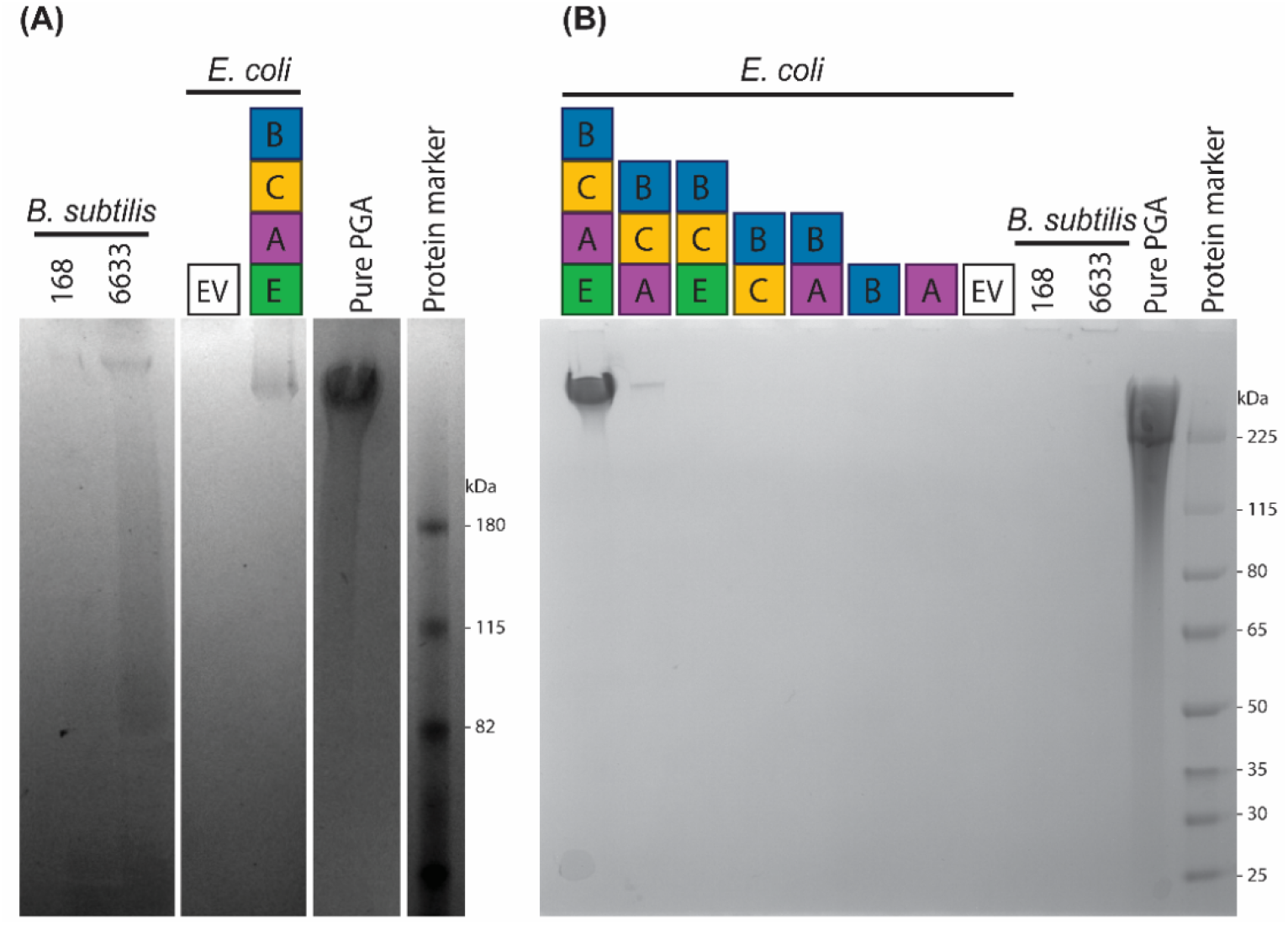
PGA produced by bacterial strains detected in cell-free culture supernatants by PAGE. (A) 48 h (“magic” medium) culture supernatants from *B. subtilis* (strain 168 and 6633) and *E. coli* MGΔΔ expressing PgsBCAE on high-copy vector. Purchased pure PGA with molecular weight > 750 kDa was used as a positive control for methylene blue staining. Rightmost lane contains a protein MW marker for reference. (B) Precipitated PGA from 24 h culture supernatant (LB medium) from *B. subtilis* and *E. coli* MGΔΔ with PgsBCAE construct on a low-copy vector. EV = empty vector negative control.

To confirm that the band detected in the PAGE corresponded to PGA, the purified samples from a PgsBCAE expressing *E. coli* culture and a commercial PGA (Sigma-Aldrich, #G1049-100MG) were hydrolyzed with HCl and analyzed by HPLC and compared to pure L-glutamate. The hydrolysate presented a single peak with a elution time similar to pure L-glutamate (Figure S2), suggesting that the polymer we purified is indeed PGA and not any other negatively-charged polymer that may be interacting with the methylene blue stain.

### 3.2. Topological prediction and expression of individual PGA synthetase subunits

The amino acid sequence of each PGS subunit was analyzed by different prediction tools in CCTOP to attest the presence of possible signal peptides and transmembrane regions. Due to the nature of each prediction algorithm and the training set that each of them uses, CCTOP compiles all results and outputs a probable topology for the sequence (Figure 2). The algorithm predicted a signal peptide for PgsB and PgsC but only presence of a transmembrane domain for PgsA and PgsE. Overall, this indicates that there is a high probability of co-localization of these proteins at cell membrane.

**Figure 2.**
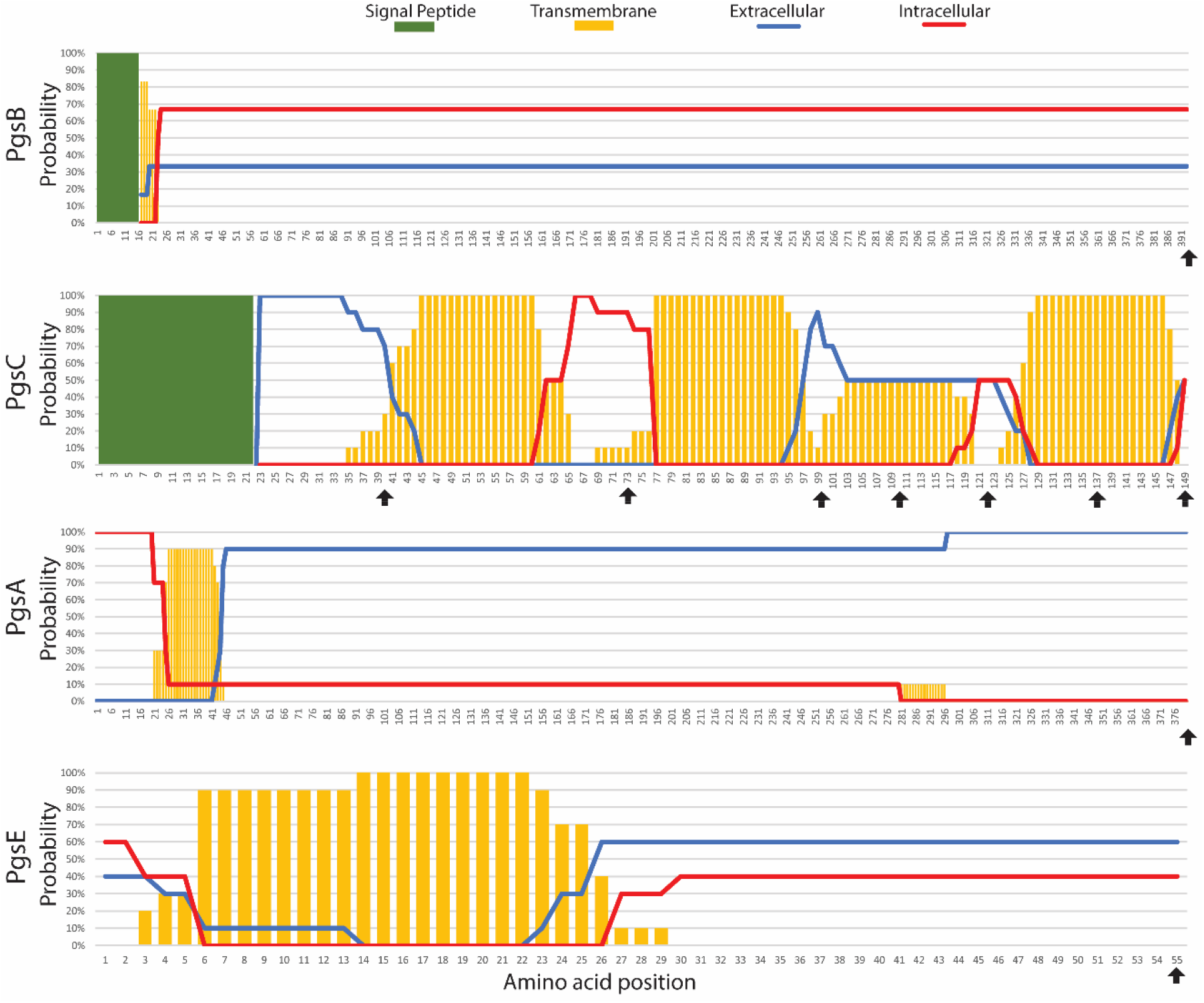
Topology prediction by CCTOP (Dobson et al., 2015). Each PGS subunit was analyzed individually by multiple prediction algorithms, and CCTOP combined the probability of each amino acid being intracellular, transmembrane, extracellular, or part of a signal peptide. Black arrows indicate residues where fusions are constructed (*vide infra*).

We also performed a created a homology model for each subunit (Figure S4). Sequence alignment by ClustalO (Sievers et al., 2011) of PgsB with *E. coli* Mur family and FolC proteins shows some conserved motifs related the ATP binding site. Construction of PgsB homology structure model with SWISS-MODEL server (Schwede et al., 2003) presented highest similarity of ~29% and sequence coverage ~80% to proteins in the Mur ligase family, which also catalyze peptide bonds (Smith, 2006). These analyses are consistent with previous studies that PgsB is homologous to other amide ligases and is the subunit responsible for catalysis (Ashiuchi et al., 2001).

To confirm the correct localization of these proteins in *E. coli*, individual PGS subunits were linked to a 6xHis tag for detection. Cells expressing tagged subunits were lysed and the soluble fractions (containing the soluble cytoplasmic proteins and small membrane fractions) and analyzed by SDS-PAGE and Western Blotting. All proteins have the expected size (PgsB: 44 kDa, PgsA: 43 kDa, PgsE: 7 kDa), except for PgsC (16 kDa), which was not detected (Figure 3). From the sequence analysis (Figure 2), we expect PgsC to have a lot of interactions with the membrane, which likely hinders proper immunodetection within the soluble fraction. Interestingly, even though PgsA and PgsE are expected to have significant transmembrane interactions, we were able to detect them in the soluble fraction, suggesting relatively weak membrane anchoring.

**Figure 3.**
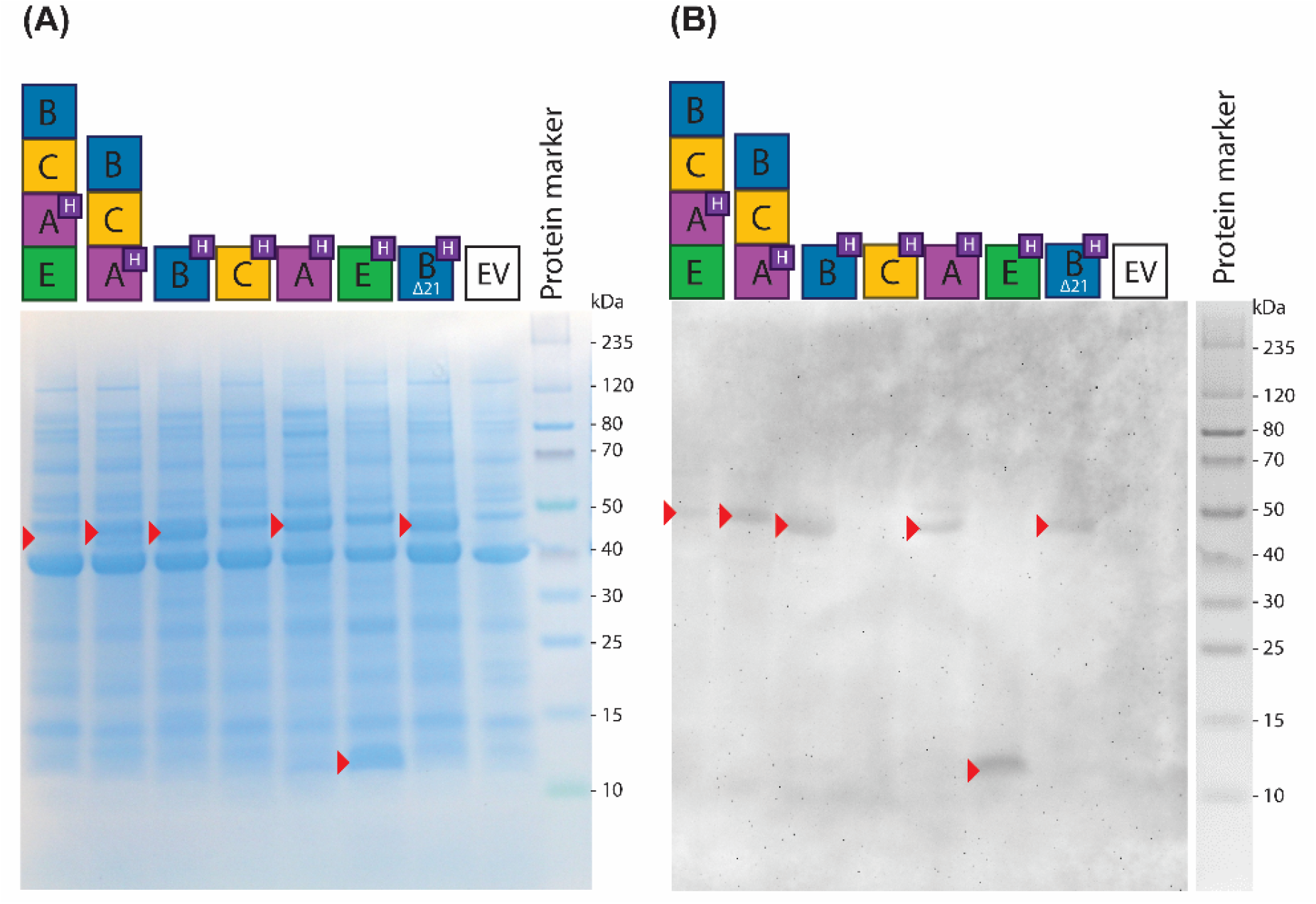
(A) SDS-PAGE of *E. coli* cell lysate expressing PgsBCAE subunits with a 6xHis-tag at the C-terminal (labeled with ‘H’ at legend). Red arrows indicate proteins present in the recombinant strains when compared to *E. coli* with empty vector control (EV). (B) Western blot of same samples using anti-6xHis antibodies.

We also prepared an additional construct of PgsB designated PgsBΔ21, which does not contain the predicted 21 amino acid N-terminal signal peptide. We expect that this variant of the protein to not be directed to the membrane and remain in the cytoplasm. We were also able to detect C-terminally tagged PgsA as part of operon structure (*pgsBCAE* and *pgsBCA*), indicating that presence of other subunits does not occlude the tag. The presence of these proteins in the soluble fraction of the lysate indicate that they are not forming inclusion bodies inside the cell, which is important to note when interpreting that the subsequent microscopy studies.

### 3.3. Localization of PGA synthetase subunits at the membrane

Fusing each subunit with a fluorescent protein (sfGFP) enables us to visualize how the different subunits localize in the cell. Superfolder GFP (sfGFP) is a highly efficient and stable folding variant, forming a folding intermediate that prevents the cysteine residues to form disulfide bonds in the periplasm (Dammeyer and Tinnefeld, 2012). Therefore, it fluoresces regardless of whether the C-terminal is exposed to the cytoplasm or periplasm. We fused each of the four subunits at their C-terminal to sfGFP and also created two variants of PgsB – one lacking the initial signal peptide (PgsBΔ21-sfGFP) and another with just the signal sequence (ssPgsB-sfGFP). The localization of full-length PgsB fusion as well as the signal peptide only fusion is evident at the membrane (Figure 4A, C), especially when compared to the N-terminally truncated (PgsBΔ21) variant (Figure 4B), where fluorescence is diffuse throughout the cytoplasm. For PgsC, PgsA, and PgsE we also note membrane localization, albeit with lower fluorescence intensities compared to PgsB (Figure 4D-F).

**Figure 4.**
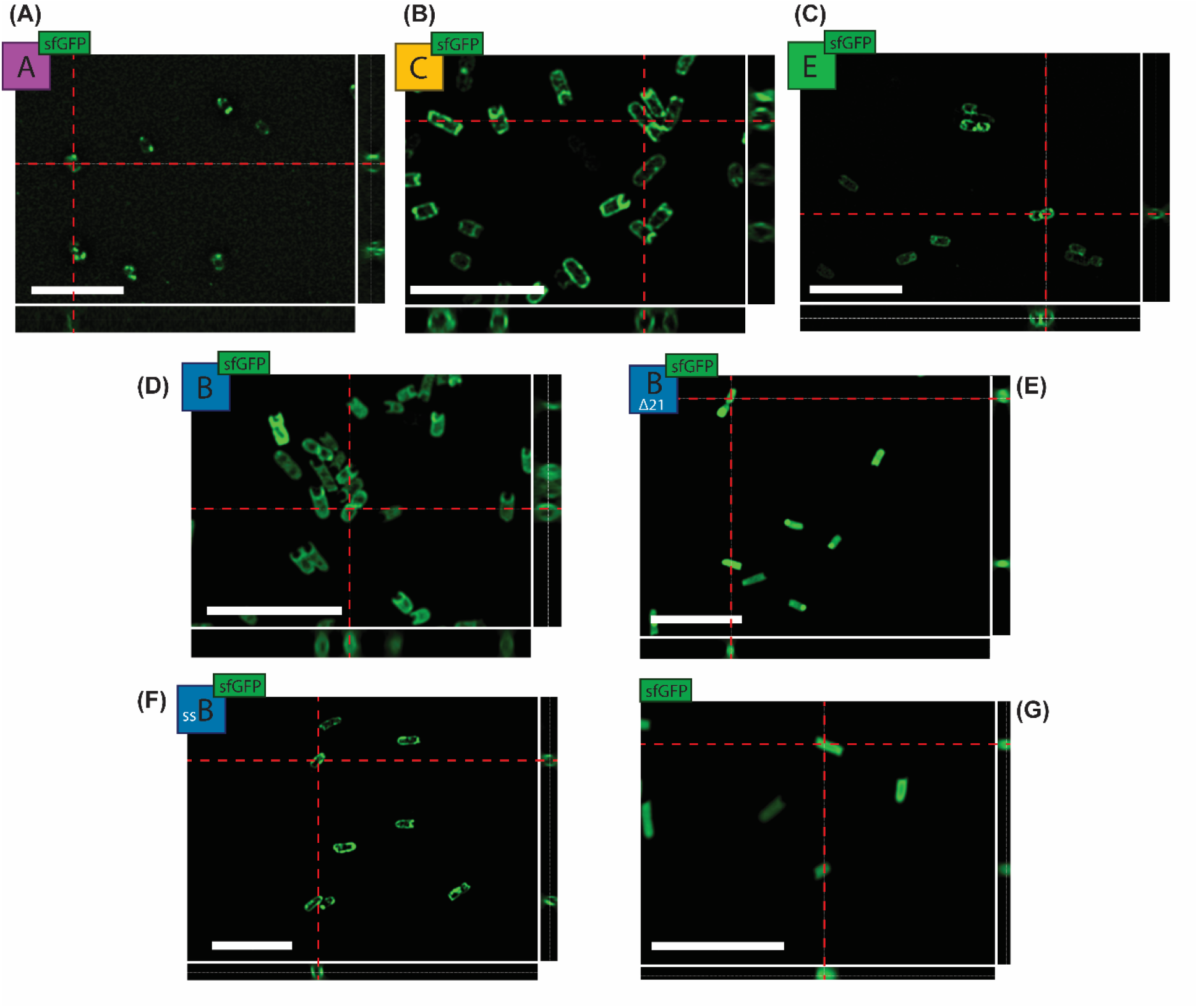
(A-D) Fluorescence microscopy of cells expressing individual subunits PgsA, C, E, and B, respectively, each with sfGFP at the C-terminus. Variants of PgsB are (E) without signal peptide (PgsBΔ21) and (F) and with only the signal peptide (ssPgsB). (G) sfGFP alone as cytoplasmic expression control. Each image represents the blind deconvolution of each captured z-stack. Side images represent orthogonal sections of the z-stack. Scale bar = 10 μm.

To determine that all components pf PGS are directed and co-localized to the *E. coli* cell membrane, we created two-fluorophore *pgs* operon constructs. All express PgsC fused to sfGFP and a second component (PgsB, A, or E) tagged with mCherry – giving PgsBrCgAE, PgsBCgArE, and PgsBCgAEr (where r = red/mCherry and g = green/sfGFP). For the PsgBCgAEr, it is evident that both proteins colocalize since we found high correlation between the green and red fluorescence signals (Figure 5). This is indicative that the proteins are correctly expressed in this heterologous host. However, for the other constructs (BrCgAE and BCgAr) we were unable to detect both fluorescent signals simultaneously, even after extensive attempts to optimize induction. We found that insertion of a fluorophore in the operon structure had a strong polarity effect on downstream genes.

**Figure 5.**
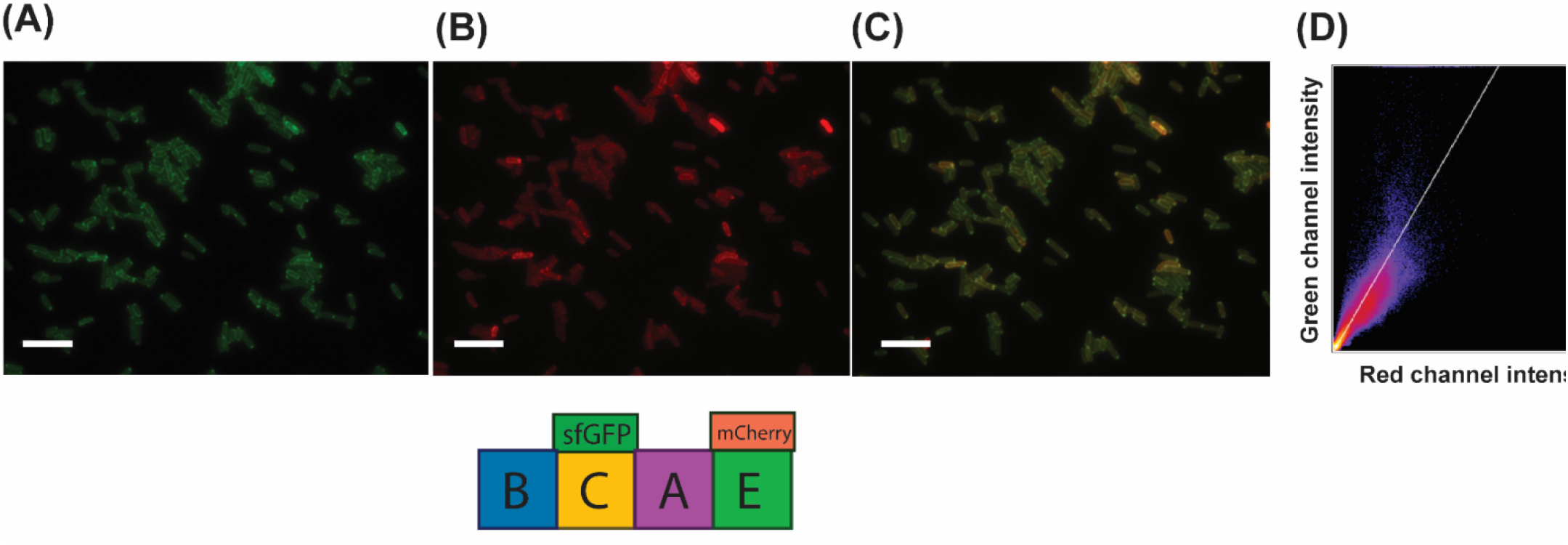
Fluorescence microscopy of cells expressing *pgsBCgAEr* operon, where the PgsC subunit is linked to sfGFP/green and PgsE to mCherry/red, both at the C-terminal. (A) GFP channel. (B) Red channel. (C) Overlay of both channels. (D) Fluorescence intensity of colocalized pixels from the green and red channels. Scale bar = 10 μm.

### 3.4. Topology of PGA synthetase subunits

Having confirmed that all four components of PGS localize to the membrane and that at least some of them co-localize, we wanted to determine the topology of each the polypeptide. The bioinformatic predictions summarized in Figure 2 (and Figure S3) indicate no clear consensus, likely due to the fact that each method uses a different training set. Therefore, we want to experimentally determine the orientation of the different subunits in the inner membrane. For this, our reporters of choice were alkaline phosphatase (PhoA) and GFPuv as C-terminal fusions. PhoA is an enzyme that only folds correctly and presents activity if exported to the periplasm of *E. coli*, where disulfide bonds can be formed. In the presence of a specific substrate such as 5-bromo-4-chloro-3-indolyl phosphate (X-Pho), colonies expression PhoA in the periplasm develop a blue color, while colonies whose enzyme is expressed in the cytoplasm remains white (Jiménez-Guerrero et al., 2013). Conversely, GFPuv only fluoresces in the cytoplasm since the oxidizing environment in the periplasm promotes the formation of disulfide bonds between C49 and C71, which causes misfolding, impeding chromophore maturation (Dammeyer and Tinnefeld, 2012).

Based on the predictions summarized in Figure 2, we expect only PgsC to have multiple transmembrane domains, with both cytoplasmic and periplasmic segments. Each of the other subunits are expected to wholly localized to either the cytoplasmic or periplasmic side of the inner membrane. We created PhoA fusions to each full-length PGS component and found that only PgsB did not develop a blue color, indicating that its C-terminus is in the cytoplasm (Figure 6A). All other components (PgsA, C, and E) developed a strong blue color signal suggesting their C-termini are periplasmically localized. For further analysis of PgsC, we created specific truncation (indicated by black arrows in Figure 2 and numbered subscripts in Figure 6) and fused them to PhoA. Figure 6A shows that only peptide 1-73 has a negative X-Pho signal, indicating that segments between 40 and 100 of PgsC are within the cytoplasm. All other truncations present as PhoA^+^, indicating periplasmic localization. To further confirm localization, we used the GFPuv fusions. Specifically, constructs that are positive for PhoA should be negative for fluorescence. In Figure 6B, PgsB presented a high signal, corroborating with the previous result that its C-terminus is in the cytoplasm. Since GFPuv fusions to full-length PgsC, PgsA, and PgsE subunits emit fluorescence that is not statistically different from the negative control (GFPuv-), we conclude that all these components terminate in the periplasmic space. While fully-folded and active GFPuv can be translocated to the periplasm by the Tat pathway (Drew et al., 2002), it is most likely that these subunits use the Sec-pathway, in which translocation is concurrent with translation. We expected, based on PhoA activity data, that PgsC fragment 1-73 would present a high fluorescence signal. However, only fragment 1-122 had fluorescence statistically different from the negative control, albeit much lower when compared the positive control (GFPuv+). Thus, the GFPuv data are not very conclusive about the topology of PgsC.

**Figure 6.**
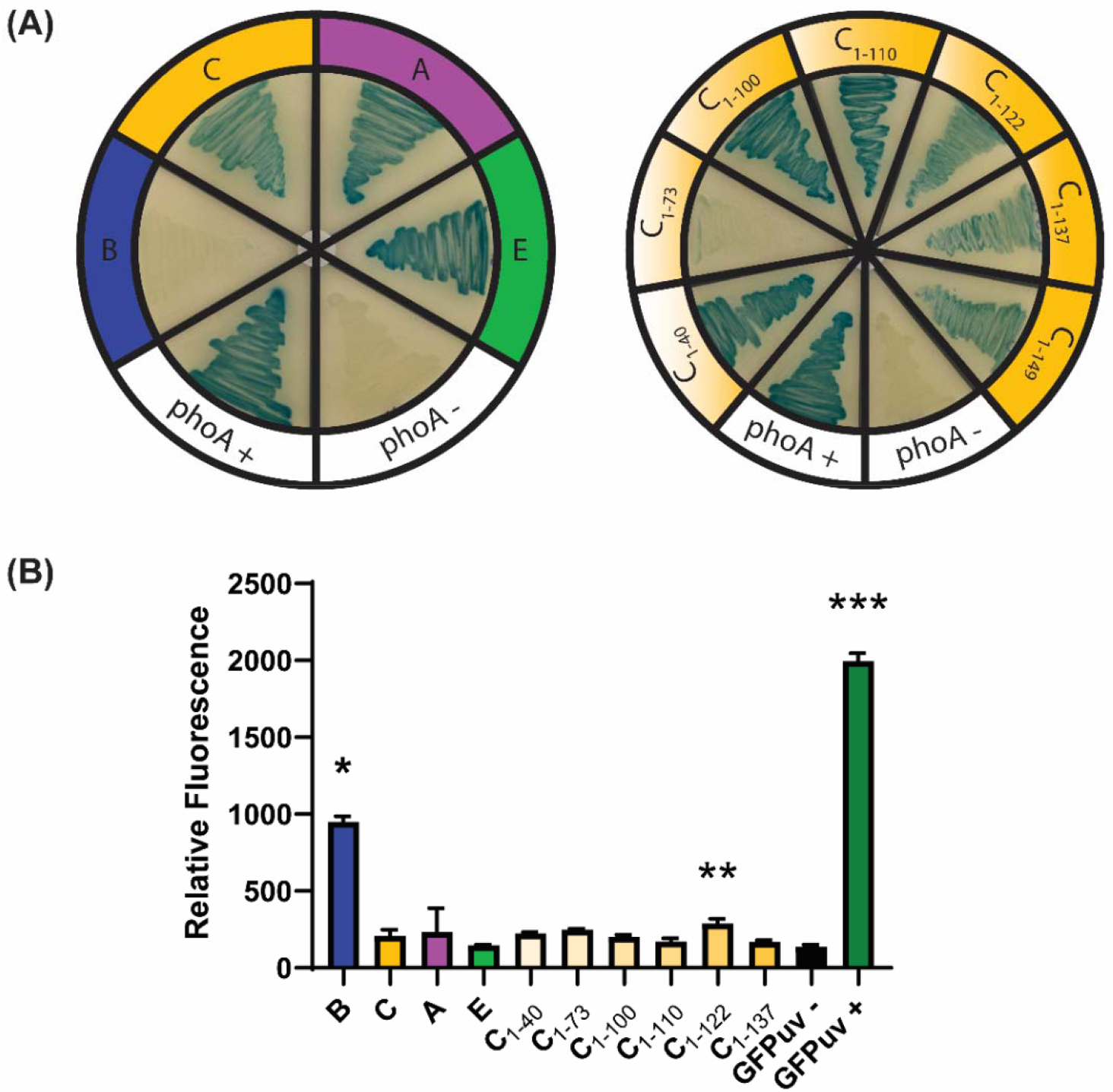
(A) LB+X-Pho plates after induction with IPTG. Each section is an *E. coli* MGΔΔ strain expressing a full-length PGS subunit with a PhoA fusion at its C-terminus. At the bottom there is a positive and negative control for PhoA activity. Truncated versions of PgsC were also assayed by the same method. (B) Relative fluorescence (RFU/OD_600_) of *E. coli* MGΔΔ strains expressing each full-length subunit with GFPuv fused at its C-terminus. Negative and positive controls constituted of cells expressing GFPuv in the periplasm and cytoplasm, respectively. Error bars indicate standard deviation of triplicate experiments. Asterisks indicate samples statistically different from negative control (p < 0.05).

Taking the PhoA and GFPuv results together, we can construct a consensus schematic of these proteins in the inner membrane of *E. coli* (Figure 7). The results for PgsB are the most conclusive, having a very distinct signal for its presence in the cytoplasm. The predicted signal peptide from amino acids 1-21 does not direct the protein to the periplasm but rather to the cytoplasmic side of the inner membrane. Based on homology predictions (Figure S4) and the proposed reaction mechanism (Ashiuchi and Misono, 2002), this subunit has access to the cytoplasmic glutamate and nucleotide pools necessary for reaction. Similarly, results were quite conclusive for PgsA and PgsE, both of which are present largely in the periplasm. Discounting the GFPuv results for PgsC, where the signals are very weak, and using only results from PhoA activity, we conclude that it has at least two (and at most three) transmembrane regions with the bulk of the protein being in the periplasm.

**Figure 7.**
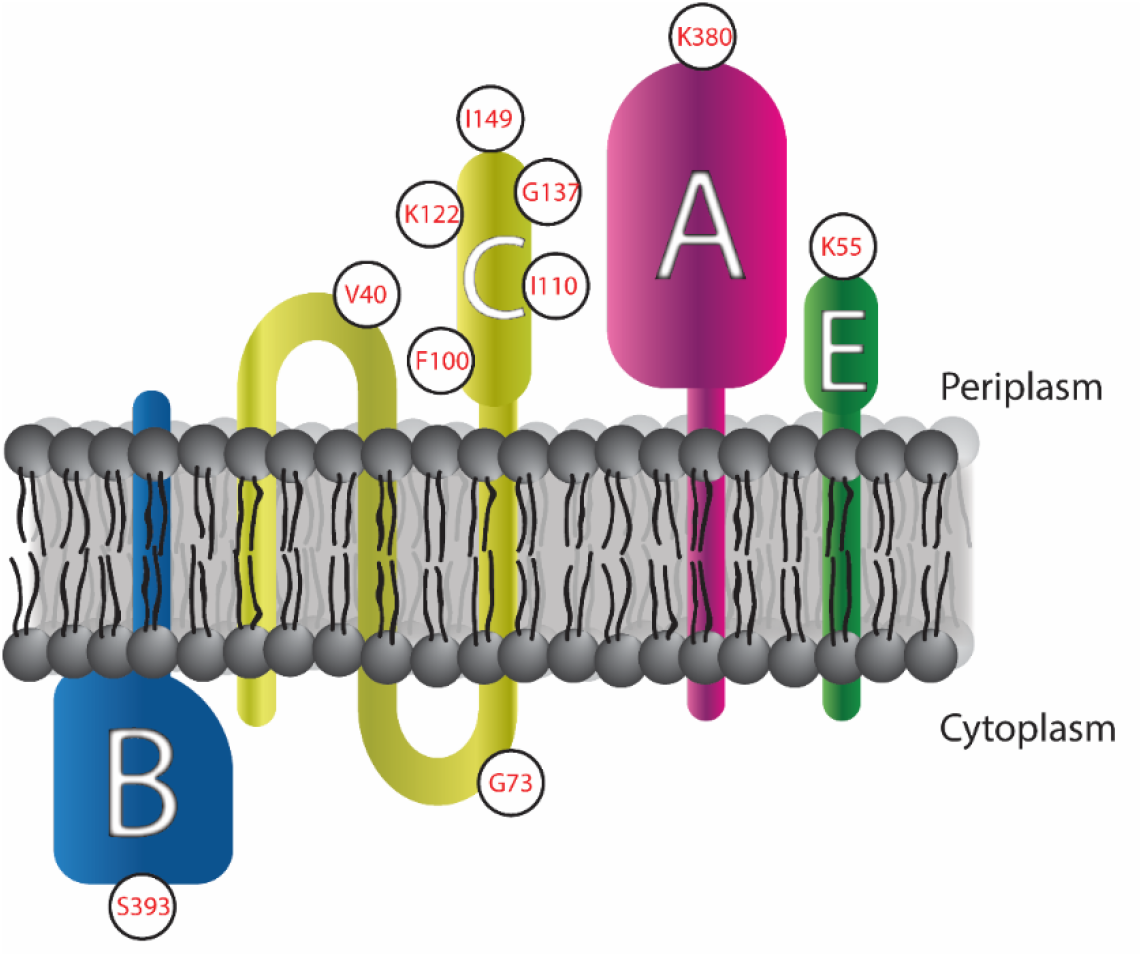
Schematic representation of PgsBCAE localization at cell membrane. White circles indicate the amino acid location where reporter fusions were added.

## 4. CONCLUSIONS

In this study, we successfully expressed the PgsBCAE membrane enzyme in a heterologous host, *E. coli*. The enzymatic complex was fully function and PGA was recovered from the supernatant of culture media. Even though production was low due to the small scale of experiments, achieving 13 mg/L of PGA, the polymer presented a high molecular weight and minimal degradation due to the absence of hydrolytic enzymes in *E. coli*. The difference in membrane structure between *B. subtilis* and *E. coli* motivated us to further investigate if the enzyme had the correct localization in the heterologous host. Not only did we observe that the tagged enzyme subunits are correctly directed to the inner cell membrane by microscopy, we could also report for the first time the orientation of the N- and C- termini of the different components across the membrane. These results will help us further understand the role of each subunit in the complex and aid in future engineering efforts.

## Abbreviations

PGA: poly-γ-glutamic acid
PGS: poly-γ-glutamic acid synthetase
MGΔΔ: *E.coli* MG1655^ΔrecAΔendA^
RFU: relative fluorescence unit
OD: optical density
MW: molecular weight
EV: empty vector

## 5. CRediT AUTHOR CONTRIBUTION STATEMENT

**Bruno Motta Nascimento:** Data curation; Formal analysis; Funding acquisition; Investigation; Methodology; Visualization; Writing - original draft & editing. **Nikhil U. Nair:** Conceptualization; Data curation; Formal analysis; Funding acquisition; Methodology; Project administration; Resources; Supervision; Visualization; Writing - review & editing.

## 7. ACKNOWLEDGEMENTS

The authors would like to thank the funding provided by NIH grant # DP2HD91798 and Tufts University (to N.U.N.), and CAPES Science without Borders grant # 13015-13-3 (to B.M.N.).

## REFERENCES

Abelson, J.N., Simon, M.I., 2009. METHODS IN ENZYMOLOGY - Guide to Protein Purification, 2nd Editio. ed. Academic Press - Elsevier. https://doi.org/10.1016/S0076-6879(09)63045-7

Ashiuchi, M., 2013. Microbial production and chemical transformation of poly-γ-glutamate. Microb. Biotechnol. 6, 664–674. https://doi.org/10.1111/1751-7915.12072

Ashiuchi, M., Misono, H., 2003. Poly-γ-glutamic acid, in: Fahnestock, S.R., Steinbüchel, A. (Eds.), Biopolymers: Volume 7 - Polyamides and Complex Proteinaceous Materials I. WILEY-VCH, pp. 123–173.

Ashiuchi, M., Misono, H., 2002. Biochemistry and molecular genetics of poly-γ-glutamate synthesis. Appl. Microbiol. Biotechnol. 59, 9–14. https://doi.org/10.1007/s00253-002-0984-x

Ashiuchi, M., Nawa, C., Kamei, T., Song, J.-J., Hong, S.-P., Sung, M.-H., Soda, K., Yagi, T., Misono, H., 2001. Physiological and biochemical characteristics of poly γ-glutamate synthetase complex of *Bacillus subtilis*. Eur. J. Biochem. 268, 5321–5328. https://doi.org/10.1046/j.0014-2956.2001.02475.x

Ashiuchi, M., Soda, K., Misono, H., 1999. A Poly-γ-glutamate Synthetic System of *Bacillus subtilis* IFO 3336: Gene Cloning and Biochemical Analysis of Poly-γ-glutamate Produced by Escherichia coli Clone Cells. Biochem. Biophys. Res. Commun. 263, 6–12.

Ashiuchi, M., Yamashiro, D., Yamamoto, K., 2013. *Bacillus subtilis* EdmS (formerly PgsE) participates in the maintenance of episomes. Plasmid 70, 209–215. https://doi.org/10.1016/j.plasmid.2013.03.008

Cao, M., Geng, W., Liu, L., Song, C., Xie, H., Guo, W., Jin, Y., Wang, S., 2011. Glutamic acid independent production of poly-γ-glutamic acid by Bacillus amyloliquefaciens LL3 and cloning of *pgsBCA* genes. Bioresour. Technol. 102, 4251–4257. https://doi.org/10.1016/j.biortech.2010.12.065

Cao, M., Geng, W., Zhang, W., Sun, J., Wang, S., Feng, J., Zheng, P., Jiang, A., Song, C., 2013. Engineering of recombinant *Escherichia coli* cells co-expressing poly-γ-glutamic acid (γ-PGA) synthetase and glutamate racemase for differential yielding of γ-PGA. Microb. Biotechnol. 6, 675–684. https://doi.org/10.1111/1751-7915.12075

Cao, M., Song, C., Jin, Y., Liu, L., Liu, J., Xie, H., Guo, W., Wang, S., 2010. Synthesis of poly (γ-glutamic acid) and heterologous expression of *pgsBCA* genes. J. Mol. Catal. B Enzym. 67, 111–116. https://doi.org/10.1016/j.molcatb.2010.07.014

Dammeyer, T., Tinnefeld, P., 2012. Engineered Fluorescent Proteins Illuminate the Bacterial Periplasm. Comput. Struct. Biotechnol. J. 3, e201210013. https://doi.org/10.5936/csbj.201210013

Datsenko, K. a, Wanner, B.L., 2000. One-step inactivation of chromosomal genes in *Escherichia coli* K-12 using PCR products. Proc. Natl. Acad. Sci. U. S. A. 97, 6640–6645. https://doi.org/10.1073/pnas.120163297

Dobson, L., Reményi, I., Tusnády, G.E., 2015. CCTOP: A Consensus Constrained TOPology prediction web server. Nucleic Acids Res. 43, W408–W412. https://doi.org/10.1093/nar/gkv451

Drew, D., Sjöstrand, D., Nilsson, J., Urbig, T., Chin, C.N., De Gier, J.W., Von Heijne, G., 2002. Rapid topology mapping of Escherichia coli inner-membrane proteins by prediction and PhoA/GFP fusion analysis. Proc. Natl. Acad. Sci. U. S. A. 99, 2690–2695. https://doi.org/10.1073/pnas.052018199

Feng, J., Gu, Y., Quan, Y., Cao, M., Gao, W., Zhang, W., Wang, S., Yang, C., Song, C., 2015. Improved poly-γ-glutamic acid production in *Bacillus amyloliquefaciens* by modular pathway engineering. Metab. Eng. 32, 106–115. https://doi.org/10.1016/j.ymben.2015.09.011

Feng, J., Quan, Y., Gu, Y., Liu, F., Huang, X., Shen, H., Dang, Y., Cao, M., Gao, W., Lu, X., Wang, Y., Song, C., Wang, S., 2017. Enhancing poly-γ-glutamic acid production in Bacillus amyloliquefaciens by introducing the glutamate synthesis features from Corynebacterium glutamicum. Microb. Cell Fact. 16, 1–12. https://doi.org/10.1186/s12934-017-0704-y

Halmschlag, B., Steurer, X., Putri, S.P., Fukusaki, E., Blank, L.M., 2019. Tailor-made poly-γ-glutamic acid production. Metab. Eng. 55, 239–248. https://doi.org/10.1016/j.ymben.2019.07.009

Hamano, Y., Arai, T., Ashiuchi, M., Kino, K., 2013. NRPSs and amide ligases producing homopoly(amino acid)s and homooligo(amino acid)s. Nat. Prod. Rep. 30, 1087–97. https://doi.org/10.1039/c3np70025a

Jiang, H., Shang, L., Yoon, S.H., Lee, S.Y., Yu, Z., 2006. Optimal production of poly-gamma-glutamic acid by metabolically engineered *Escherichia coli*. Biotechnol. Lett. 28, 1241–1246. https://doi.org/10.1007/s10529-006-9080-0

Jiménez-Guerrero, I., Cubo, M., Pérez-Montaño, F., López-Baena, F., Guash-Vidal, B., Ollero, F., Bellogín, R., Espuny, M., 2013. Bacterial Protein Secretion Systems, Beneficial Plant-microbial Interactions. https://doi.org/10.1201/b15251-10

Karimova, G., Ladant, D., 2017. Defining Membrane Protein Topology Using *pho-lac* Reporter Fusions, in: Journet, L., Cascales, E. (Eds.), Bacterial Protein Secretion Systems. Methods in Molecular Biology, Vol 1615. Humana Press, New York, NY, pp. 129–142.

Ke, N., Landgraf, D., Paulsson, J., Berkmen, M., 2016. Visualization of periplasmic and cytoplasmic proteins with a self-labeling protein tag. J. Bacteriol. 198, 1035–1043. https://doi.org/10.1128/JB.00864-15

Kino, K., Arai, T., Arimura, Y., 2011. Poly-α-glutamic acid synthesis using a novel catalytic activity of RimK from Escherichia coli K-12. Appl. Environ. Microbiol. 77, 2019–2025. https://doi.org/10.1128/AEM.02043-10

Liu, C.L., Dong, H.G., Xue, K., Yang, W., Liu, P., Cai, D., Liu, X., Yang, Y., Bai, Z., 2019. Biosynthesis of poly-γ-glutamic acid in *Escherichia coli* by heterologous expression of pgsBCAE operon from *Bacillus*. J. Appl. Microbiol. 128, 1390–1399. https://doi.org/10.1111/jam.14552

Manocha, B., Margaritis, A., 2010. A novel method for the selective recovery and purification of γ-polyglutamic acid from *Bacillus licheniformis* fermentation broth. Biotechnol. Prog. 26, 734–742. https://doi.org/10.1002/btpr.370

Ogunleye, A., Bhat, A., Irorere, V.U., Hill, D., Williams, C., Radecka, I., 2015. Poly-γ-glutamic acid: production, properties and applications. Microbiology 161, 1–17. https://doi.org/10.1099/mic.0.081448-0

Ojima, Y., Kobayashi, J., Doi, T., Azuma, M., 2019. Knockout of *pgdS* and ggt gene changes poly-γ-glutamic acid production in Bacillus licheniformis RK14-46. J. Biotechnol. 304, 57–62. https://doi.org/10.1016/j.jbiotec.2019.08.003

Park, C., Choi, J.C., Choi, Y.H., Nakamura, H., Shimanouchi, K., Horiuchi, T., Misono, H., Sewaki, T., Soda, K., Ashiuchi, M., Sung, M.H., 2005. Synthesis of super-high-molecular-weight poly-γ-glutamic acid by *Bacillus subtilis* subsp. chungkookjang. J. Mol. Catal. B Enzym. 35, 128–133. https://doi.org/10.1016/j.molcatb.2005.06.007

Sambrook, J., Russell, D.W., 2001. Molecular Cloning - A Laboratory Manual - Vol. 1, 2 and 3, Third Edit. ed. Cold Spring Harbor Laboratory Press, New York.

Schwede, T., Kopp, J., Guex, N., Peitsch, M.C., 2003. SWISS-MODEL: An automated protein homology-modeling server. Nucleic Acids Res. 31, 3381–3385. https://doi.org/10.1093/nar/gkg520

Scoffone, V., Dondi, D., Biino, G., Borghese, G., Pasini, D., Galizzi, A., Calvio, C., 2013. Knockout of *pgdS* and *ggt* genes improves γ-PGA yield in B. subtilis. Biotechnol. Bioeng. 110, 2006–2012. https://doi.org/10.1002/bit.24846

Sievers, F., Wilm, A., Dineen, D., Gibson, T.J., Karplus, K., Li, W., Lopez, R., McWilliam, H., Remmert, M., Söding, J., Thompson, J.D., Higgins, D.G., 2011. Fast, scalable generation of high-quality protein multiple sequence alignments using Clustal Omega. Mol. Syst. Biol. 7. https://doi.org/10.1038/msb.2011.75

Smith, C.A., 2006. Structure, Function and Dynamics in the mur Family of Bacterial Cell Wall Ligases. J. Mol. Biol. 362, 640–655. https://doi.org/10.1016/j.jmb.2006.07.066

Soto, A.M., Draper, D., 2012. White gels: An easy way to preserve methylene blue stained gels. Anal. Biochem. 421, 345–346. https://doi.org/10.1016/j.ab.2011.10.048

Tarui, Y., Iida, H., Ono, E., Miki, W., Hirasawa, E., Fujita, K.I., Tanaka, T., Taniguchi, M., 2005. Biosynthesis of poly-γ-glutamic acid in plants: Transient expression of poly-γ-glutamate synthetase complex in tobacco leaves. J. Biosci. Bioeng. 100, 443–448. https://doi.org/10.1263/jbb.100.443

Tian, G., Fu, J., Wei, X., Ji, Z., Ma, X., Qi, G., Chen, S., 2014. Enhanced expression of *pgdS* gene for high production of poly-γ-glutamic aicd with lower molecular weight in *Bacillus licheniformis* WX-02. J. Chem. Technol. Biotechnol. 89, 1825–1832. https://doi.org/10.1002/jctb.4261

Wang, N., Yang, G., Che, C., Liu, Y., 2011. Heterogenous expression of poly-gamma-glutamic acid synthetase complex gene of *Bacillus licheniformis* WBL-3. Appl. Biochem. Microbiol. 47, 381–385. https://doi.org/10.1134/s0003683811040193

Wang, Q., Wei, X., Chen, S., 2017. Production and Application of Poly-γ-glutamic Acid, Current Developments in Biotechnology and Bioengineering. Elsevier B.V. https://doi.org/10.1016/B978-0-444-63662-1.00030-0

Xavier, J.R., Madhan Kumarr, M.M., Natarajan, G., Ramana, K.V., Semwal, A.D., 2019. Optimized production of poly (γ-glutamic acid) (γ-PGA) using Bacillus licheniformis and its application as cryoprotectant for probiotics. Biotechnol. Appl. Biochem. 1–12. https://doi.org/10.1002/bab.1879

Xu, G., Zha, J., Cheng, H., Ibrahim, M.H.A., Yang, F., Dalton, H., Cao, R., Zhu, Y., Fang, J., Chi, K., Zheng, P., Zhang, X., Shi, J., Xu, Z., Gross, R.A., Koffas, M.A.G., 2019. Engineering *Corynebacterium glutamicum* for the de novo biosynthesis of tailored poly-γ-glutamic acid. Metab. Eng. 56, 39–49. https://doi.org/10.1016/j.ymben.2019.08.011

Yamashiro, D., Minouchi, Y., Ashiuchi, M., 2011a. Moonlighting role of a poly-γ-glutamate synthetase component from *Bacillus subtilis*: insight into novel extrachromosomal DNA maintenance. Appl. Environ. Microbiol. 77, 2796–2798. https://doi.org/10.1128/AEM.02649-10

Yamashiro, D., Yoshioka, M., Ashiuchi, M., 2011b. Bacillus subtilis *pgsE* (formerly *ywtC*) stimulates poly-γ-glutamate production in the presence of zinc. Biotechnol. Bioeng. 108, 226–230. https://doi.org/10.1002/bit.22913

Yoon, S.H., Do, J.H., Lee, S.Y., Chang, H.N., 2000. Production of poly-γ -glutamic acid by fed-batch culture of *Bacillus licheniformis*. Time 585–588.

Yuan, Z., Ran, Q., Chang, Z., Gao, H., Jia, C., 2019. Recovery of low-molecular-weight γ-PGA by metal cation from the fermentation broth. Process Biochem. 82, 215–221. https://doi.org/10.1016/j.procbio.2019.04.001

