## Supplemental Information for "Characterization of a membrane enzymatic complex for heterologous production of poly-γ-glutamate in *E. coli*"

Table S1 – Bacterial strains used in this study.

| **Strain** | | **Description** | **Source** |
| --- | --- | --- | --- |
| *Escherichia coli* | NEB5α | fhuA2 Δ(*argF-lacZ*)U169 *phoA* *glnV44* Φ80 Δ(*lacZ*)M15 *gyrA96* *recA1* *relA1* *endA1* *thi*-1 *hsdR17* | New England Biolabs |
|  | MG1655(DE3)^ΔrecA,ΔendA^ (MG∆∆) | *E. coli* str. K-12 F– λ– *ilvG*– *rfb*-50 *rph*-1 λ(DE3) *ΔrecA* *ΔendA* | This work |
|  | MG∆∆-∆phoA | MG1655(DE3)^ΔrecA,ΔendA^ *ΔphoA* | This work |
| *Bacillus subtilis* | ATCC 6633 | *B. subtilis* (Ehrenberg) Cohn | USDA-NRRL |
|  | 168 | *B. subtilis* Marburg mutagenesis, *trpC2* | USDA-NRRL |

Table S2 – Plasmids used in this study.

| **Plasmids** | **Description** | **Source** |
| --- | --- | --- |
| pACYC-Duet1 | Double multiple cloning sites with T7 promoter/terminator, chloramphenicol resistance (*cat*) | EMDMillipore |
| pET-Duet1 | Double multiple cloning sites with T7 promoter/terminator, ampicillin resistance (*bla*) | EMDMillipore |
| pACYC-BCAE | pACYC-Duet1 backbone with *pgsBCAE* operon | This work |
| pACYC-BCA | pACYC-Duet1 backbone with *pgsBCA* operon | This work |
| pACYC-BCE | pACYC-Duet1 backbone with *pgsBCE* operon | This work |
| pET-BCA | pET-Duet1 backbone with *pgsBCA* operon | This work |
| pET-BCE | pET-Duet1 backbone with *pgsBCE* operon | This work |
| pET-CAE | pET-Duet1 backbone with *pgsCAE* operon | This work |
| pET-BAE | pET-Duet1 backbone with *pgsBAE* operon | This work |
| pACYC-BC | pACYC-Duet1 backbone with *pgsBC* operon | This work |
| pACYC-BA | pACYC-Duet1 backbone with *pgsBA* operon | This work |
| pACYC-B | pACYC-Duet1 backbone with *pgsB* gene | This work |
| pACYC-C | pACYC-Duet1 backbone with *pgsC* gene | This work |
| pACYC-A | pACYC-Duet1 backbone with *pgsA* gene | This work |
| pACYC-E | pACYC-Duet1 backbone with *pgsE* gene | This work |
| pXXX-His | pACYC-Duet1 backbone, XXX= described operon or gene, 6xHis tag | This work |
| pXXX-mCherry | pACYC-Duet1 backbone, XXX= described operon or gene, mCherry tag | This work |
| pXXX-sfGFP | pACYC-Duet1 backbone, XXX= described operon or gene, sfGFP tag | This work |
| pXXX-GFPuv | pACYC-Duet1 backbone, XXX= described operon or gene, GFPuv tag | This work |
| pXXX-phoA | pACYC-Duet1 backbone, XXX= described operon or gene, phoA tag | This work |
| pKD46 | ampicillin resistance (*bla*), lambda red recombinase (exo, β, γ), temperature-sensitive replication (repA101ts) | (Datsenko and Wanner, 2000) |
| pKD3 | chloramphenicol resistance (*cat*), FRT sites | (Datsenko and Wanner, 2000) |
| pCP20 | ampicillin resistance (*bla*), chloramphenicol resistance (*cat*), temperature-sensitive replication (repA101ts), thermal induction of FLP recombinase | (Datsenko and Wanner, 2000) |

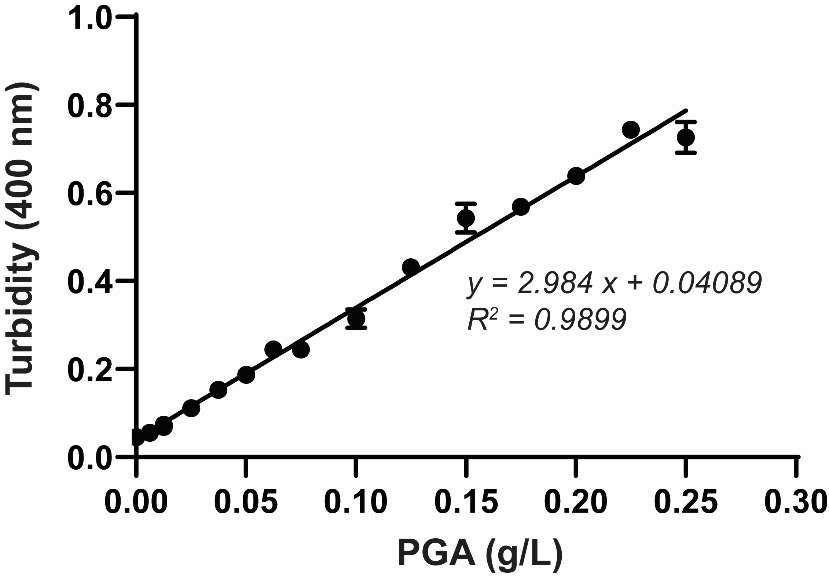

Fig. S1 – Cetyl trimethyammonium bromide (CTAB) assay standard curve with pure PGA.

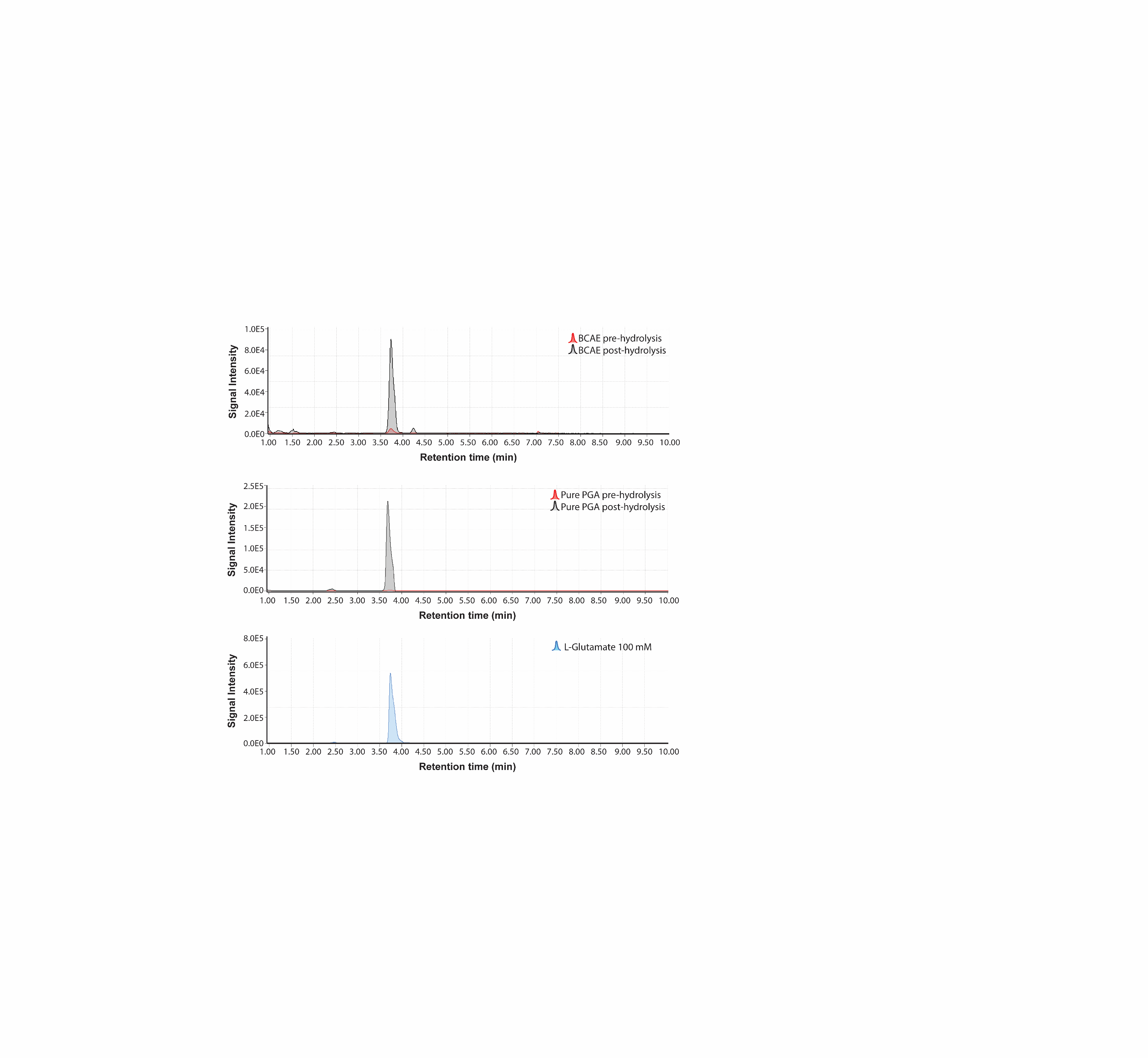

(C)

(B)

(A)

Fig. S2 – HPLC of pre- and post-hydrolysis samples from (A) purified PGA from *E. coli* MG∆∆ PgsBCAE culture, (B) pure commercial PGA, and (C) pure L-glutamate.

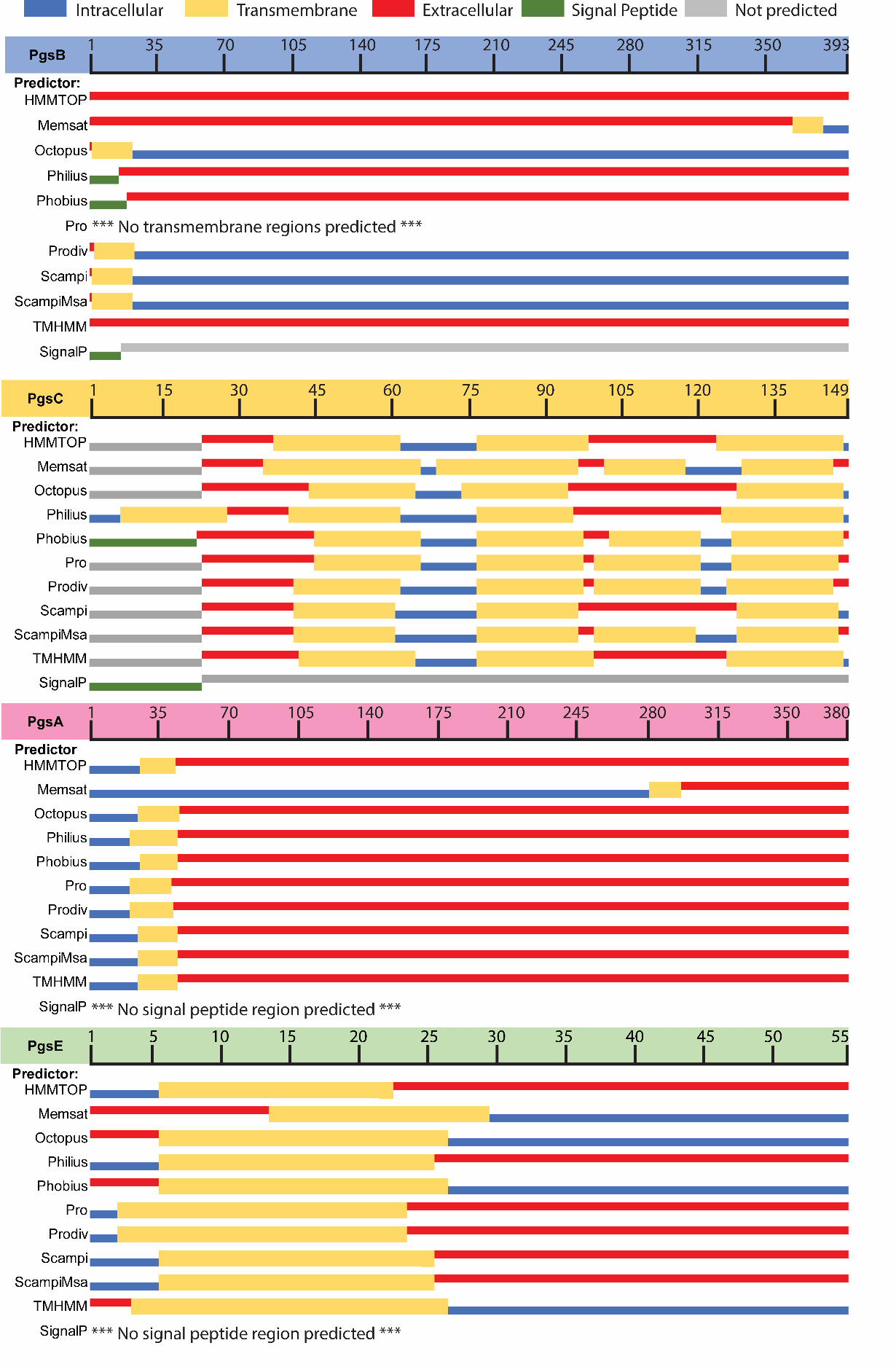

Fig. S3 – Individual topology prediction by different server algorithms (HMMTOP, Memsat, Octopus, Philius, Phobius, Pro, Prodiv, Scampi-single, Scampi-msa, TMHMM, SignalP) (Dobson et al., 2015).

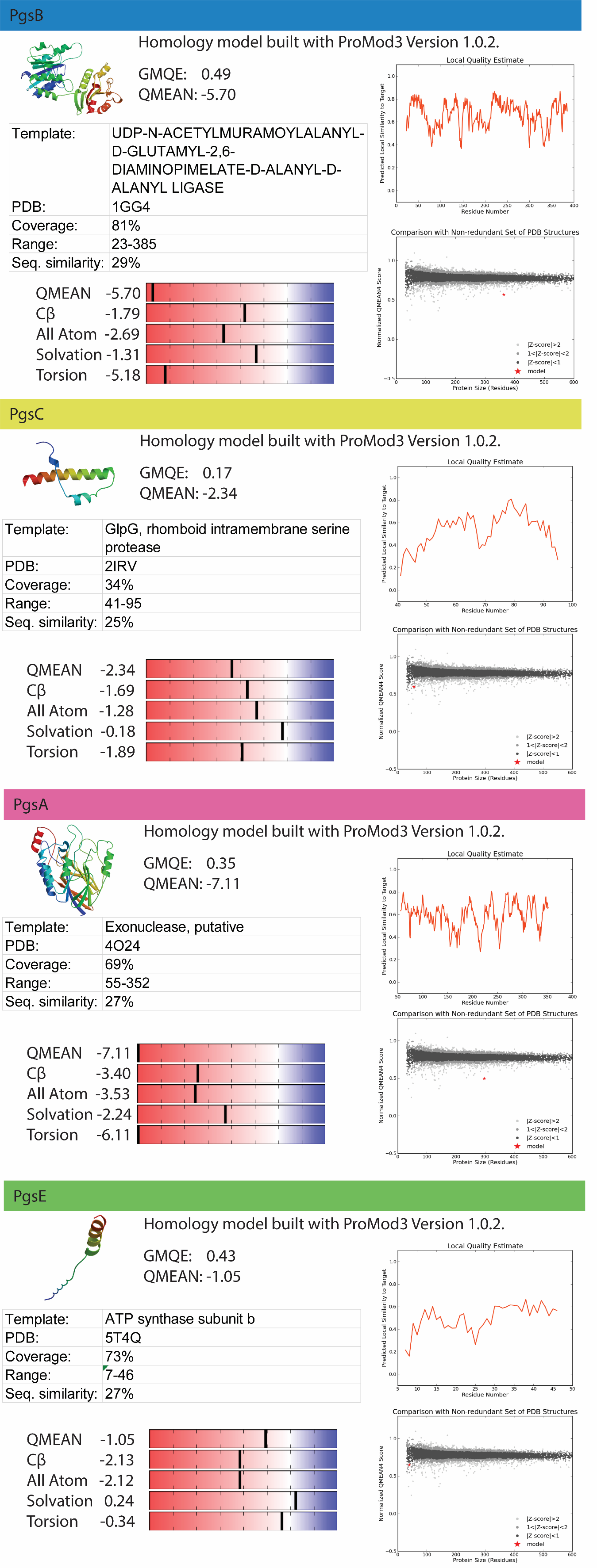

Fig. S4 – Homology models for each PgsBCAE subunit obtained with SWISS-MODEL server (Schwede et al., 2003).

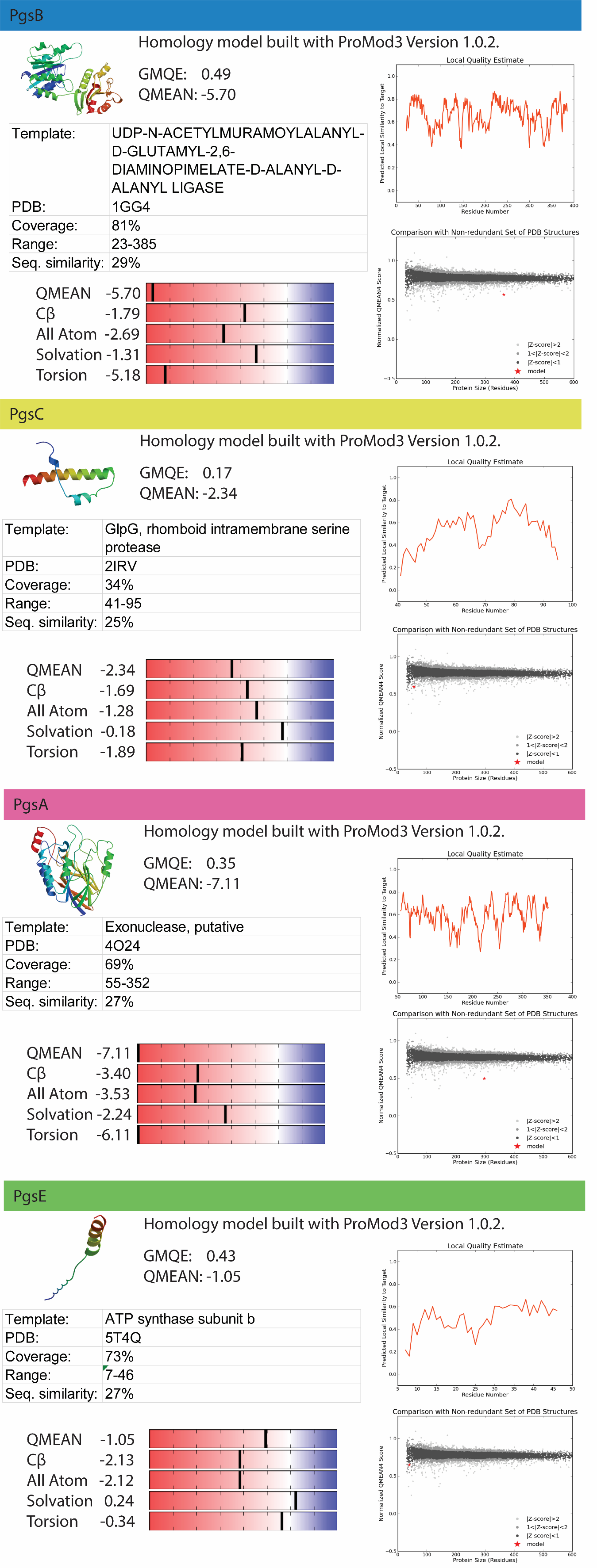

Fig. S4 (cont.) – Homology models for each PgsBCAE subunit obtained with SWISS-MODEL server (Schwede et al., 2003).

**REFERENCES**

Datsenko, K. a, Wanner, B.L., 2000. One-step inactivation of chromosomal genes in *Escherichia coli* K-12 using PCR products. Proc. Natl. Acad. Sci. U. S. A. 97, 6640–6645. https://doi.org/10.1073/pnas.120163297

Dobson, L., Reményi, I., Tusnády, G.E., 2015. CCTOP: A Consensus Constrained TOPology prediction web server. Nucleic Acids Res. 43, W408–W412. https://doi.org/10.1093/nar/gkv451

Schwede, T., Kopp, J., Guex, N., Peitsch, M.C., 2003. SWISS-MODEL: An automated protein homology-modeling server. Nucleic Acids Res. 31, 3381–3385. https://doi.org/10.1093/nar/gkg520
